# Including random effects in statistical models in ecology: fewer than five levels?

**DOI:** 10.1101/2021.04.11.439357

**Authors:** Dylan G.E. Gomes

## Abstract

As generalized linear mixed-effects models (GLMMs) have become a widespread tool in ecology, the need to guide the use of such tools is increasingly important. One common guideline is that one needs at least five levels of a random effect. Having such few levels makes the estimation of the variance of random effects terms (such as ecological sites, individuals, or populations) difficult, but it need not muddy one’s ability to estimate fixed effects terms – which are often of primary interest in ecology. Here, I simulate ecological datasets and fit simple models and show that having too few random effects terms does not influence the parameter estimates or uncertainty around those estimates for fixed effects terms. Thus, it should be acceptable to use fewer levels of random effects if one is not interested in making inference about the random effects terms (i.e. they are ‘nuisance’ parameters used to group non-independent data). I also use simulations to assess the potential for pseudoreplication in (generalized) linear models (LMs), when random effects are explicitly ignored and find that LMs do not show increased type-I errors compared to their mixed-effects model counterparts. Instead, LM uncertainty (and p values) appears to be more conservative in an analysis with a real ecological dataset presented here. These results challenge the view that it is never appropriate to model random effects terms with fewer than five levels – specifically when inference is not being made for the random effects, but suggest that in simple cases LMs might be robust to ignored random effects terms. Given the widespread accessibility of GLMMs in ecology and evolution, future simulation studies and further assessments of these statistical methods are necessary to understand the consequences of both violating and blindly following simple guidelines.

## Introduction

As ecological datasets are inherently messy and researchers gain increased access to data, statistical analyses in ecology are becoming more complex (Low-Décarie, Chivers & Granados, 2014), and advances in computing power and freely available statistical software are increasing the accessibility of such analyses to non-statisticians (Bates et al., 2007; Patil, Huard & Fonnesbeck, 2010; Gabry & Goodrich, 2016; Salvatier, Wiecki & Fonnesbeck, 2016; Bürkner, 2017; Carpenter et al., 2017; Magnusson et al., 2017; Rue et al., 2017). As these methods have become more complex and accessible to ecologists, fisheries and wildlife managers, and evolutionary biologists the need to guide the use of such tools is becoming increasingly important (Bolker, 2008; Bolker et al., 2009; Zuur, Ieno & Elphick, 2010; Kéry & Royle, 2015; Kass et al., 2016; Zuur & Ieno, 2016; Harrison et al., 2018; Arnqvist, 2020; Silk, Harrison & Hodgson, 2020). The use of generalized linear mixed-effects models (GLMM), for example, has become a widespread tool that allows one to build hierarchical models that can estimate, and thus account for, imperfect detection in biological surveys (e.g. occupancy, N-mixture, mark-recapture, etc.) and can model correlations among data that come from non-independent groups or populations (i.e. random effects; also known as varying effects) (Bolker, 2008; Kéry & Royle, 2015; Powell & Gale, 2015; Harrison et al., 2018; McElreath, 2020).

Generalized linear mixed-effects models are a regression type analysis that are flexible in that they can handle a variety of data generating processes such as binomial (e.g. presence / absence of a species, alive / dead individuals) and Poisson (e.g. wildlife counts). When the sampling distribution is Gaussian (also known as normal; e.g. mean-centered continuous data such as: tree diameter, vocalization frequency, or body condition residuals), this is a special case of a GLMM that is referred to as simply a linear mixed-effects model (LMM). GLMMs (and LMMs) differ from their simpler counterparts, (generalized) linear models (GLMs and LMs), in that they include random effects, in addition to the fixed effects (hence *mixed-effects*).

Fixed effects are also often called predictors, covariates, explanatory or independent variables in ecology and include both variables of interest (e.g. *average temperature* in climate change studies or *sound pressure levels* in anthropogenic noise research) and other variables that are only included to control for unexplained variation but often not directly useful to understanding the research question at hand (e.g. date or time of sampling in the above studies). Fixed effects are *fixed* in that the model parameters (*β* in equation 1 below) are assumed to be fixed, or non-random, and are not drawn from a hypothetical distribution.

Random effects, on the other hand, allow one to combine information (e.g. in a meta-analysis), deal with spatiotemporal autocorrelation, use partial pooling to borrow strength from other populations or groups, account for grouping or blocked designs (e.g. repeat-measures data from sites or individuals), and estimate population-level parameters, among others (Kéry & Royle, 2015). Thus, the random effects structure should be decided by the experimental design (Barr et al., 2013; Arnqvist, 2020). Random effects are *random* in that they are assumed to be drawn randomly from a distribution – often a Gaussian distribution – during the data-generating process. This is most often done by fitting a random *intercept* for each group (see equation 1 below), but one can, and should, also assign random *slopes* to variables, where the slopes of variables (not just the intercepts) are allowed to vary by group (see Bolker, 2008; Schielzeth & Forstmeier, 2009; Kéry & Royle, 2015; Harrison et al., 2018). If we are interested in the variability of a population (of individuals, groups, sites, or populations), it is difficult to estimate this variation with too few levels of individuals, groups, sites, or populations (i.e. random effects terms).

> “When the number of groups is small (less than five, say), there is typically not enough information to accurately estimate group-level variation” (Gelman & Hill, 2006).
>
> “…if interest lies in measuring the variation among random effects, a certain number is required…To obtain an adequate estimate of the among-population heterogeneity – that is, the variance parameter – at least 5 - 10 populations might be required” (Kéry & Royle, 2015).
>
> “With <5 levels, the mixed model may not be able to estimate the among-population variance accurately” (Harrison et al., 2018).
>
> “Strive to have a reasonable number of levels (at the very least, say, four to five subjects) of your random effects within each group” (Arnqvist, 2020).

This guideline that random effects terms should have at least five levels (i.e. groups) is backed by only limited empirical evidence (Harrison, 2015), but it is intuitive that too few draws from distribution will hinder one’s ability to estimate the variance of that distribution. Indeed, in each of the above segments of quoted text, the authors suggest that five levels are needed for *estimation of group-level, or among-population, variance*. However, this rule is often adhered to out of context, where authors or reviewers of ecological journals suggest that one cannot use random effects terms if they do not contain at least five levels, *in any case*.

Simulations by Harrison (2015) demonstrate that random effects variance can be biased more strongly when the levels of random effects terms are low, yet in this work it appears that slope (beta) estimates for fixed effects terms are generally not more biased with only three random effects levels compared to five. There are many cases (and some would argue that in *most cases*, see below) in which the variance of random effects is not directly of interest to the ecological research question at hand.

> “…in the vast majority of examples of random-effects (or mixed) models in ecology, the random effects do *not* have a clear ecological interpretation. Rather, they are merely abstract constructs invoked to explain the fact that some measurements are more similar to each other than others are – i .e., to model correlations in the observed data” (Kéry & Royle, 2015).

Thus, it is unclear whether or not it is appropriate to use random effects when there are fewer than five grouping levels in situations where one does not directly care about the ‘nuisance’ among-population variance, but instead is interested in estimates and uncertainty of predictor variables (i.e. fixed effects). The current state of practice in ecology is to drop the random effects terms such that we are now using generalized linear models where we are not grouping observations (we drop the **M**ixed-effects from the GL**M**M to become GLM). I question whether we are choosing to accept pseudoreplication in ecology (Hurlbert, 1984; Kéry & Royle, 2015; Arnqvist, 2020), rather than inaccurate estimates of among-population variance. In cases where one does not care about among-population variance, this tradeoff may be non-existent, but little research exists to support this.

Here, I simulate ecological datasets to assess whether *fixed effects* estimates are more biased when the accompanying *random effects* consist of fewer than five levels; I also ask whether using an alternative model without random effects (LMs) leads to higher type I errors (demonstrating a ‘significant’ effect when in fact one does not exist). I also analyze a real dataset within a similar framework to understand how the number of random effects levels and model structure (random effects or not) might change ecological inference.

### Methodology

All simulation of datasets and model fitting was done in R v4.0.4 (R Core Team, 2017), all visualizations were accomplished with the aid of R package ‘ggplot2’ (Wickham, 2011).

#### Data generation

I used a modified version of code from Harrison (2015), to explore the importance of varying two parameters in a linear mixed-effect model (LMM): the number of observations in a dataset (30, 60, or 120), and the number of levels of the random intercept term (3, 5, or 20). We can think of the latter as the number of individuals in an experiment or the number of field sites in a study. This was done by generating a response variable *y*_*i*_ from the following equation:

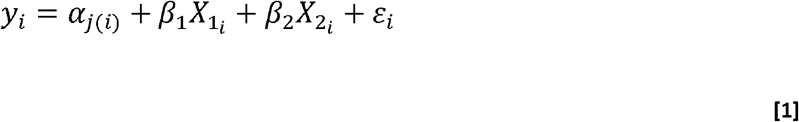

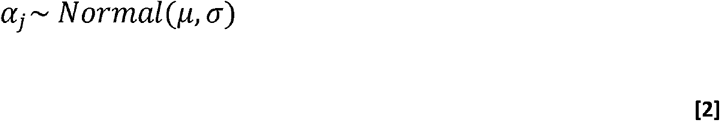

Where *α*_*j(i)*_ is the intercept for site (or individual) *j* to which observation *i* belongs. Thus, each observation shared a site-level intercept, which were drawn from a normal distribution with mean (μ) = 0 and standard deviation (σ) = 0.5. *β*_1_ and *β*_2_ are the slope parameters for two generic predictor variables (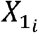 and 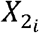 respectively), which were both randomly generated from a normal distribution with μ = 0 and σ = 0.5, which mimics standardized variables that are centered by their mean and scaled by two standard deviations (Gelman, 2008). Note that while 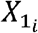 and 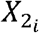 were drawn from a normal distribution during data generation, their associated parameters *β*_1_ and *β*_2_ are not (these are the *fixed* effects). For all simulated datasets, parameter values were fixed at *β*_1_ = 2 and *β*_2_ = 0, meaning 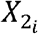 does not have a linear relationship with, or is only randomly related to, the response variable y_i_. This allows for an assessment of type-I error rate, since any significant *p* values for this *β*_2_ slope parameter are erroneous. The error term ε_i_ is unique to each observation *i* that is drawn from a normal distribution with µ_ε_= 0 and σ_ε_ = 0.25 (same as equation 2 above); this term is simply adding noise to our system during the data generating process, but is modelled implicitly as residual variance in many generic GLMM functions.

As an ecological example, you might think of the above equations as

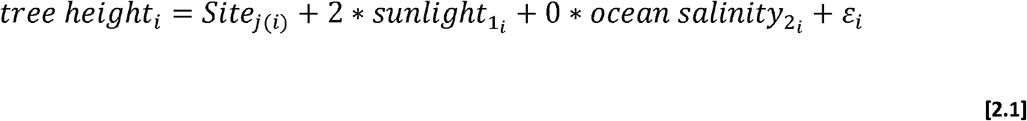

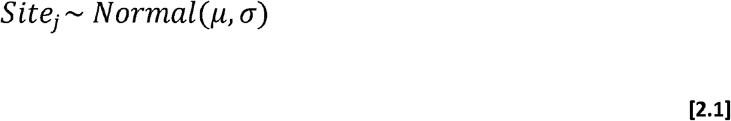

Data from each site or forest patch (Site_1_, Site_2_, …, Site_n_) are grouped by their own site-level intercept, which is randomly drawn from a normal distribution; some sites have shorter trees on average and other sites have taller trees on average, but the entire forest (where all sites were randomly selected from; equation 2.1) has a mean height of µ and a variance σ. We might expect sunlight to have a positive relationship with tree height (*β*_1_ = 2) and ocean salinity to reasonably have no relationship with tree height (*β*_2_ = 0). We could just leave the ocean salinity term out of the equation entirely, but this explicitly demonstrates that the parameter coefficient that we estimate with the model should = 0.

#### Model fitting simulations

For each of the nine combinations of scenarios (30, 60, or 120 observations by 3, 5, or 20 sites), I simulated 10,000 datasets. Each dataset was fit with a linear mixed-effect model (LMM) and a linear model (LM). All model fitting was done with R functions ‘lmer’ (LMM) or ‘lm’ (LM) in the package ‘lme4’ or in ‘base’ R, respectively (Bates et al., 2007; R Core Team, 2017), and p values for ‘lme4’ models were calculated with ‘lmerTest’ (Kuznetsova, Brockhoff & Christensen, 2017), see note in discussion about this.

~~~
#LMM:
m1 <-lmer(y ∼ x_1_ + x_2_ + (1|Site))
m1 <-lmer(tree.height ∼ sunlight + ocean.salinity + (1|Site))
R Code
~~~

Where x_1_ and x_2_ are fixed effects (see equation 1), and `(1|Site)` is the syntax for specifying a random intercept (*α*_*j(i)*_ in equation 1). In ecology, we often fit independent sites as unique levels of a random effect, so I use site here for demonstration purposes. But site can be replaced with individual, group, population, etc. Often the recommendation, if one has fewer than 5 levels of random effects terms (*j* < 5 in *α*_*j(i)*_), is to fit the random effects as fixed effects (LMM becomes LM), specified in R as:

~~~
#LM:
m2 <-lm(y ∼ x_1_ + x_2_ + Site)
m2 <-lm(tree.height ∼ sunlight + ocean.salinity + Site)
R Code
~~~

and mathematically defined as:

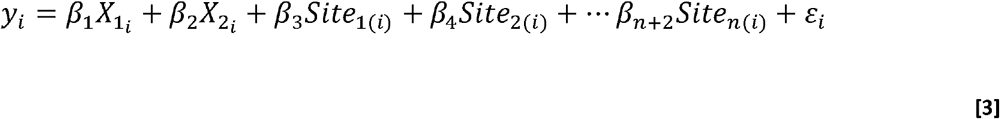

Now a *β* term is estimated for each site (or population) level independently. Site effects no longer come from a normal distribution (as in equation 2), but instead are considered fixed, hence *fixed effects*. Thus, both a LMM and a LM were fit to each simulated dataset (n = 10,000) of each of the nine combinations of data-generation (30, 60, or 120 observations by 3, 5, or 20 sites; 90,000 total simulated datasets and 180,000 models fit to data). This allowed for a comparison of the type-I error rates of LMMs and LMs, the latter of which ignores the blocked structure of data (i.e. site-level grouping).

#### Type-I error calculation and p values

Type-I error rate was calculated as the proportion of 10,000 models that a ‘significant’ p value of ≤ 0.05 was obtained for the *β*_2_ parameter estimate (e.g. ocean salinity) in which the true value of that parameter was set to be 0. I sampled (with replacement) 10,000 p value ‘observations’ from each group of 10,000 models to produce a new proportion of type-I error; this process was repeated 1,000 times, and the bootstrapped 95% confidence intervals were calculated as the 0.025 and 0.975 quantiles of those 1,000 replications (see code; modified from Harrison, 2015).

#### Case study: spider body condition and noise

I used orb-weaving spider body condition data from a previous experiment (Gomes, Hesselberg & Barber, 2021). In short, speakers were used to experimentally broadcast whitewater river noise (predictor variable: sound pressure level) over the course of multiple summers. Orb-weaving spiders (*Tetragnatha versicolor*) were then weighed and measured (femur length), and body condition (response variable) was calculated from a residual index, using these measurements (Jakob, Marshall & Uetz, 1996; Gomes, 2020).

Here, I randomly sampled this dataset to create 30,000 new datasets, such that each new dataset contained three random observations from each of 3, 5, or 10 sites (10,000 datasets for each level of random effects terms). Thus, each dataset contained 9, 15, or 30 total observations when there were 3, 5, or 10 sites, respectively. While it would have been ideal to separate sample size from the number of random effects (as in the simulated datasets above), this simply wasn’t possible to do with this real dataset (i.e. obtaining 30 total observations from three sites was not possible, while easily done for ten sites). Each dataset was fit with a linear mixed-effect model (LMM) and a linear model (LM) for comparison. Similar to the simulations above, all model fitting was done with R functions ‘lmer’ (LMM) or ‘lm’ (LM) in the package ‘lme4’ or in ‘base’ R, respectively (Bates et al., 2007; R Core Team, 2017) with the formulae:

~~~
m3 <-lmer(body.condition ∼ sound.pressure.level + (1|Site)) # LMM
m4 <-lm(body.condition ∼ sound.pressure.level + Site) # LM
                                                                                              R Code
~~~

I used the same methodology as above in ‘*Type-I error calculation*’ to calculate the proportion of 10,000 models in which *p* < 0.05, which corresponds to rejecting the null hypothesis that sound pressure level is not related to spider body condition. In this case, we are not assessing a type-I error, because we do not truly know if this variable should be significant or not, but instead it serves as a point of comparisons across methods (LMM vs LM) with real data rather than simulated.

## Results

### Estimating model parameters and uncertainty

Linear mixed models and linear models were able to resurrect simulated fixed effect relationships with no noticeable patterns in bias, regardless of number of levels of random effects or sample size. That is, both mean model parameter estimates (*β*_1_ and *β*_2_) were centered on their true values (2 and 0, respectively; Table 1; Figure 1, S1). The uncertainty around these estimates generally decreased as sample size increased. For example, doubling the sample size from 30 observations to 60 observations lead to a decrease by 36.6% and 35.5% in parameter estimate uncertainty (for *β*_1_ and *β*_2_ respectively; Table 1; Figure 1). Another doubling to 120 observations lead to a further decrease in uncertainty by 33.4% and 32.9%, respectively. The number of levels of random effects appears to be relatively non-important in resurrecting model parameter estimates within these simulation scenarios (Table 1; Figure 1); instead there were small, likely negligible, increases in uncertainty around fixed effect parameter estimates as the number of levels of random effects increased.

**Table 1:**
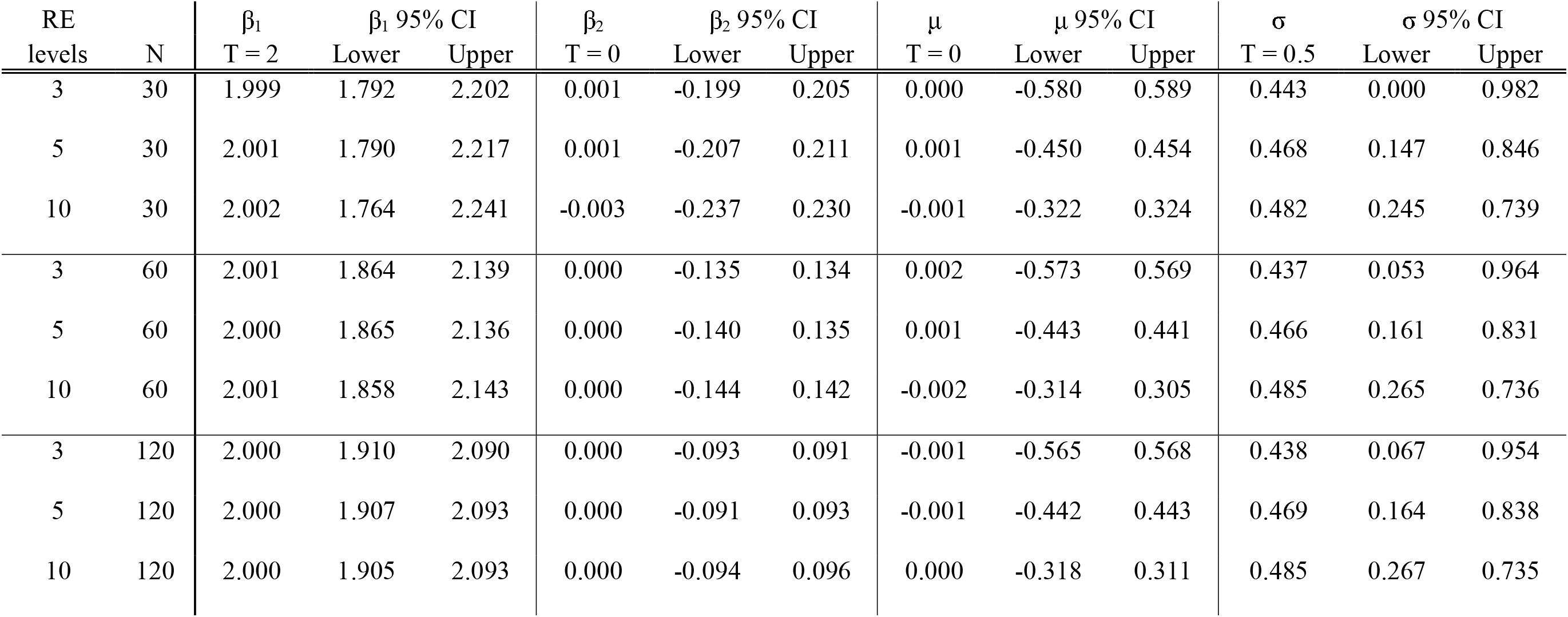
Model estimates from 10,000 simulated datasets. The number of levels of random effects (RE) was varied (3, 5, or 10), as was the number of observations in the dataset (N = 30, 60, or 120). The true (T) values for the data generation process (equation 1) are indicated in the second header row underneath the estimated parameter labels (fixed effects: β_1_, β_2_; random effects: μ, σ). The mean of 10,000 model estimates (β_1_, β_2_, μ, σ) are indicated for the respective models below the true values. Lower and upper bounds on 95% confidence intervals for each parameter is calculated as the 0.025 and 0.975 quantiles, respectively, of 1,000 bootstrapped replications (see methods).

**Figure 1:**
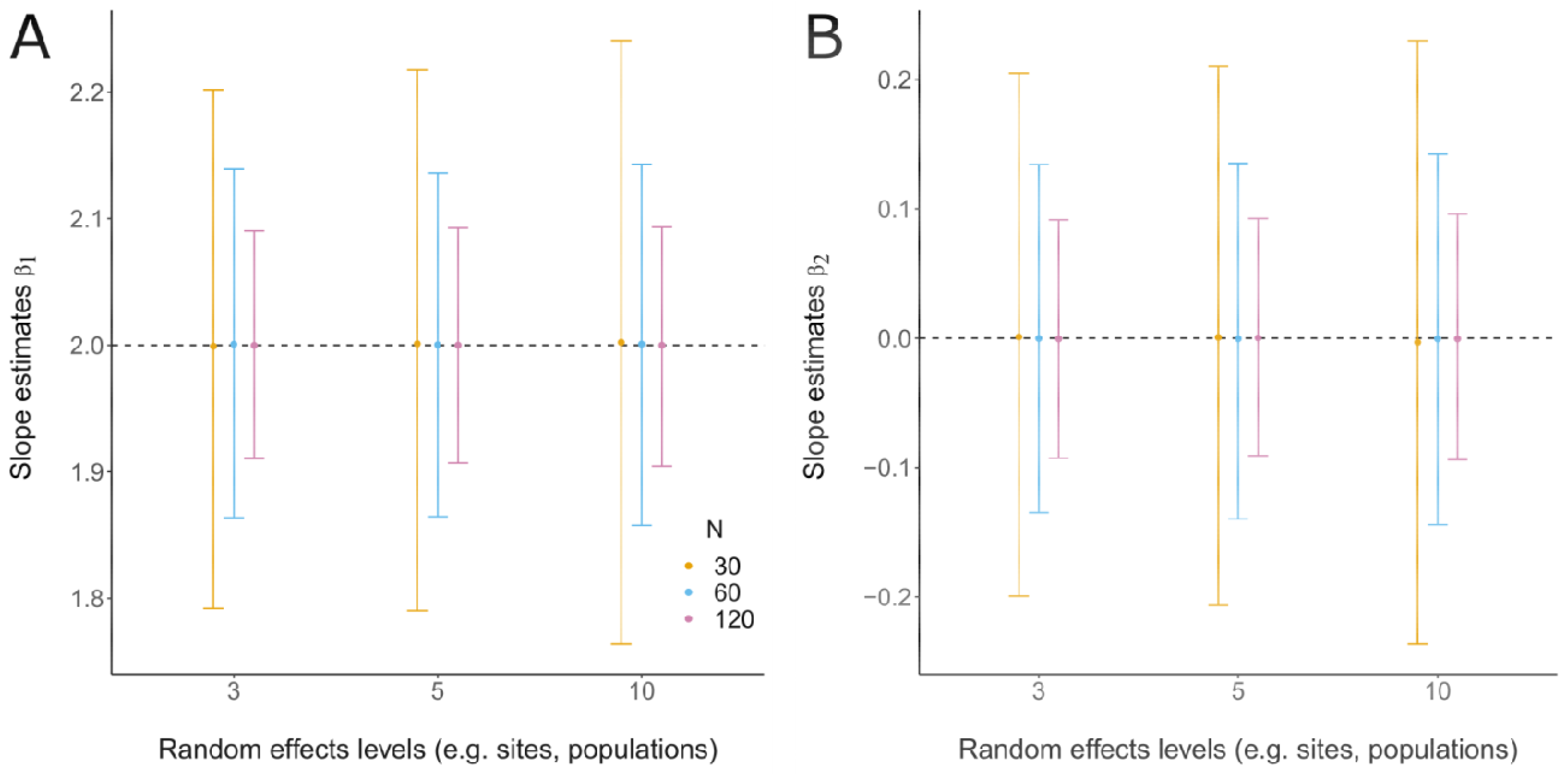
Fixed effects model estimates for simulated data. Each point is the mean estimate for 10,000 models (and datasets), whereas error bars are 95% confidence intervals. N = the number of observations (i.e. number of rows) in each dataset. Dashed lines indicate the true value. In all scenarios the bias in parameter estimates are negligible. As the sample size increases, our certainty around the parameter estimates (β) increases, but the number of random effects has a relatively minor effect on estimating β. When sample sizes (N) are low, parameter uncertainty increases with increasing levels of random effects (assuming a consistent N).

All LMM estimates of the distribution mean (μ) were unbiased, regardless of number of levels of random effects or sample size (Table 1; Figure 2A). The random effects variance (σ) estimates, however, were not centered at the true value, and estimates were more biased with fewer levels of random effects, whereas sample size did not affect this bias (Table 1; Figure 2B). That is, with only three levels of random effects the magnitude of the bias was 12.2% of the true value. Increasing to five levels of random effects nearly halved this bias to 6.4%, and increasing to 10 levels halved the bias again to 3.2% of the true value. Averaged across numbers of random effects terms, estimates were biased by about 7% regardless of sample size (7.1%, 7.4%, and 7.2% for N = 30, 60, and 120 respectively).

**Figure 2:**
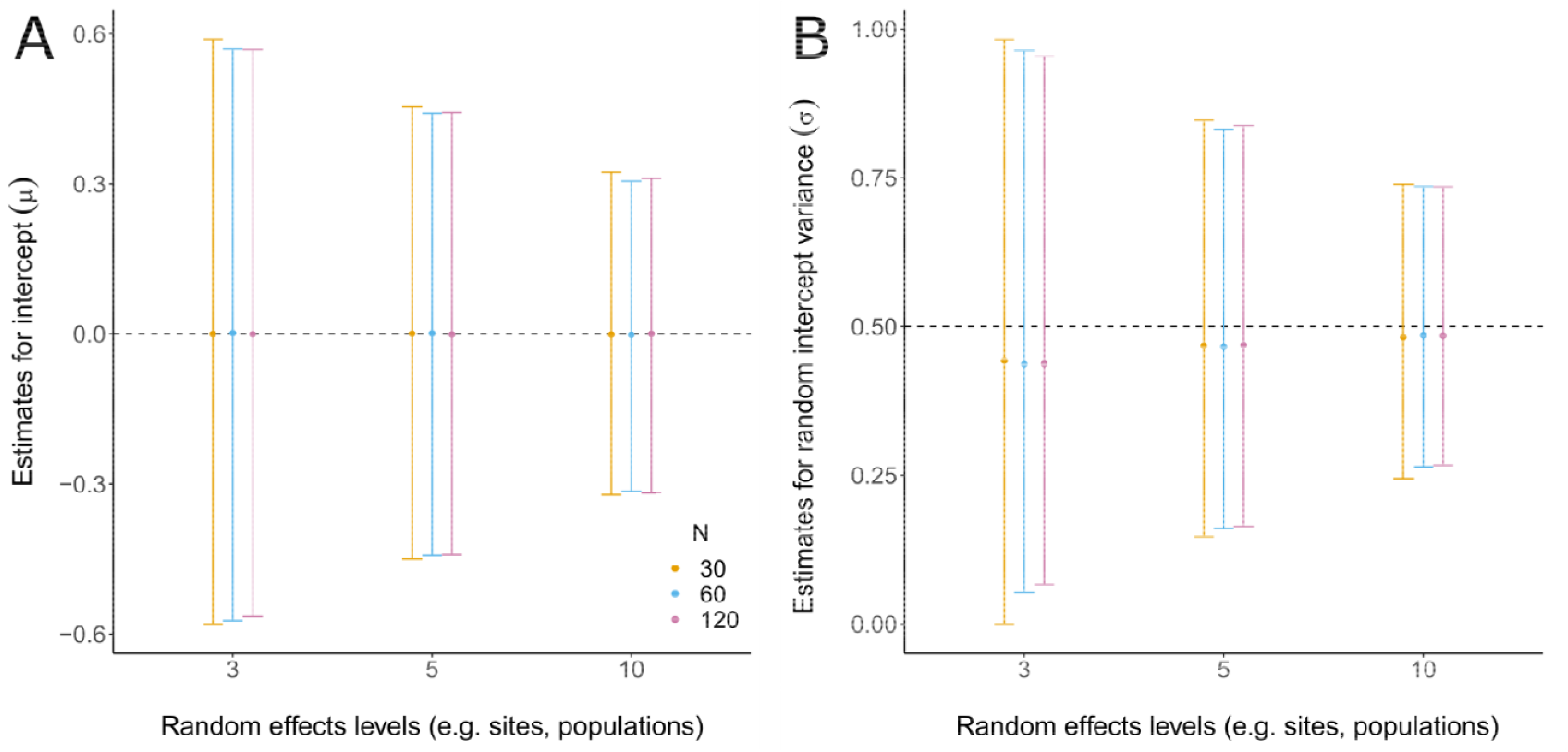
Random effects model estimates for simulated data. Each point is the mean estimate for 10,000 models (and datasets), whereas error bars are 95% confidence intervals. N = the number of observations (i.e. number of rows) in each dataset. Dashed lines indicate the true value. A) As the number of random effects levels increases, the uncertainty around the mean (μ) decreases. Sample size has a relatively minor effect on estimating μ. B) As the number of random effects levels increases, the bias and uncertainty around the random effects variance (σ) decreases. Sample size has a small, but relatively minor effect on estimating σ. The bias in σ starts to approach the starting (simulated) σ = 0.5 as the number of random effects reaches 10.

The uncertainty around random effects estimates (μ and σ) generally decreased with an increased number of random effects levels, whereas sample size did little to alleviate this uncertainty (Table 1; Figure 2). Increasing the number of random effects levels from 3 to 5, and then from 5 to 10, decreased the uncertainty for μ by 22.4% and 29.1%, respectively, and for σ by 26.6% and 29.8% respectively.

#### Type-I errors

For all simulated datasets, both LMM and LM produced type-I error rates around the typical α = 0.05, with 95% confidence intervals overlapping this value. Neither sample size, nor the number of random effects levels seemed to influence the type-I error rate. Furthermore, dropping the random effects structure (using a LM instead of a LMM) did not increase the probability type-I errors (Figure 3), nor the ability of the model to accurately estimate fixed effects parameters (fixed effects estimates appear the same as they do in Figure 1, see Figure S1).

**Figure 3:**
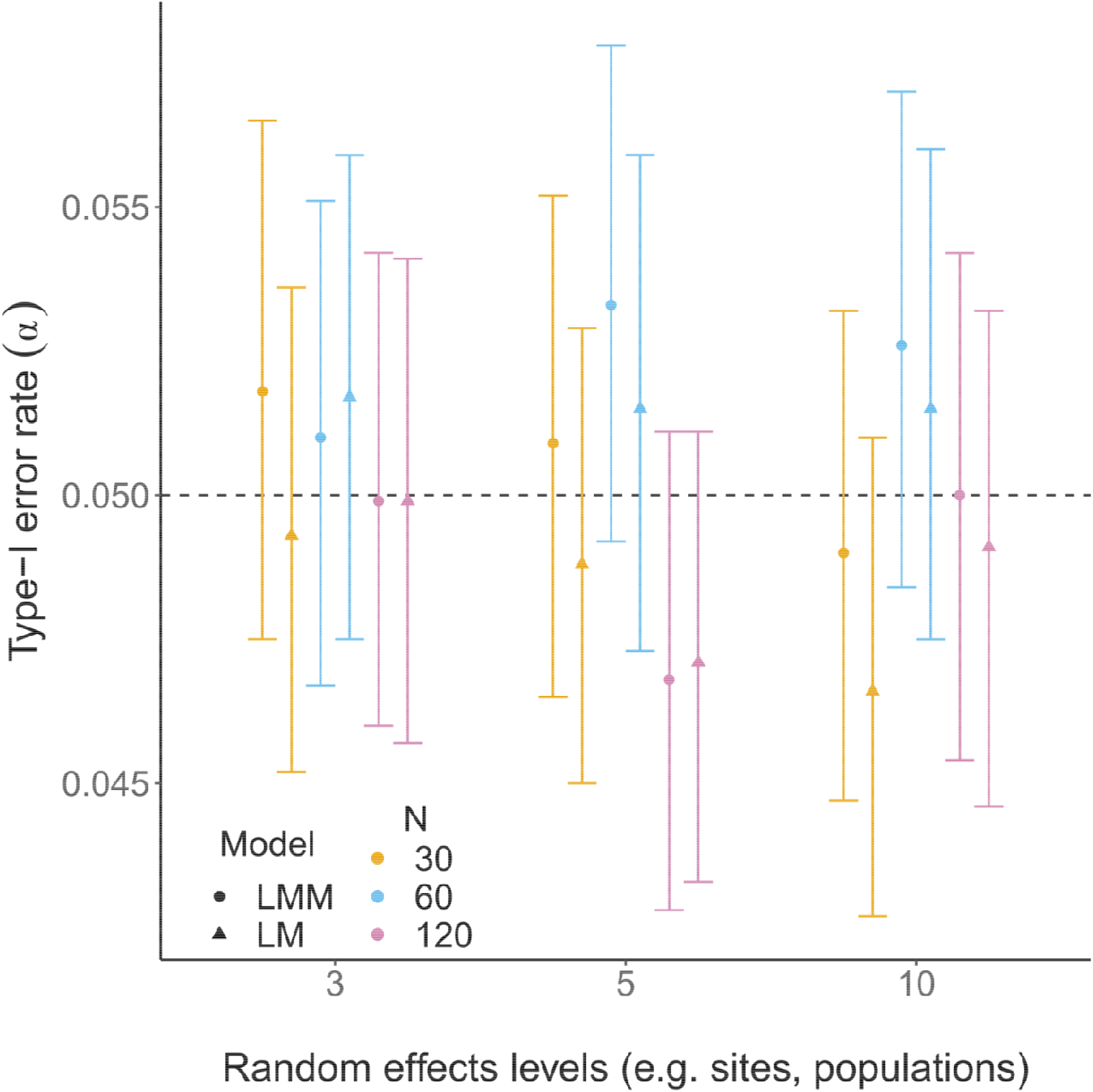
Type-I error for various linear models (LM) and linear mixed-effects models (LMM). Type-I error rate was calculated as the proportion of models (n = 10,000) in which a ‘significant’ p value of ≤ 0.05 was obtained for a parameter estimate in which the true value of that parameter was set to be 0 (Figure 1B); each point represents this proportion. To generate error bars as 95% confidence intervals, I used bootstrapping to replicate this process 1,000 times (see methods). N = the number of observations (i.e. number of rows) in each dataset. Symbols indicate model type (LM vs LMM). Dashed lines indicate the true alpha value (0.05).

#### Case study results

Linear mixed models (LMM) and linear models (LM) both consistently estimated the fixed effect parameters for sound pressure level to be very weakly positive, with large overlap with zero (or no relationship). Linear model estimates are slightly shrunk towards zero, compared to LMMs (Figure 4A), but it is impossible to assess bias, since we do not know truth with these biological data. The uncertainty around these estimates decreases with LMMs, compared to LMs, and decreases as the sample size and number of sites increases (in tandem).

**Figure 4:**
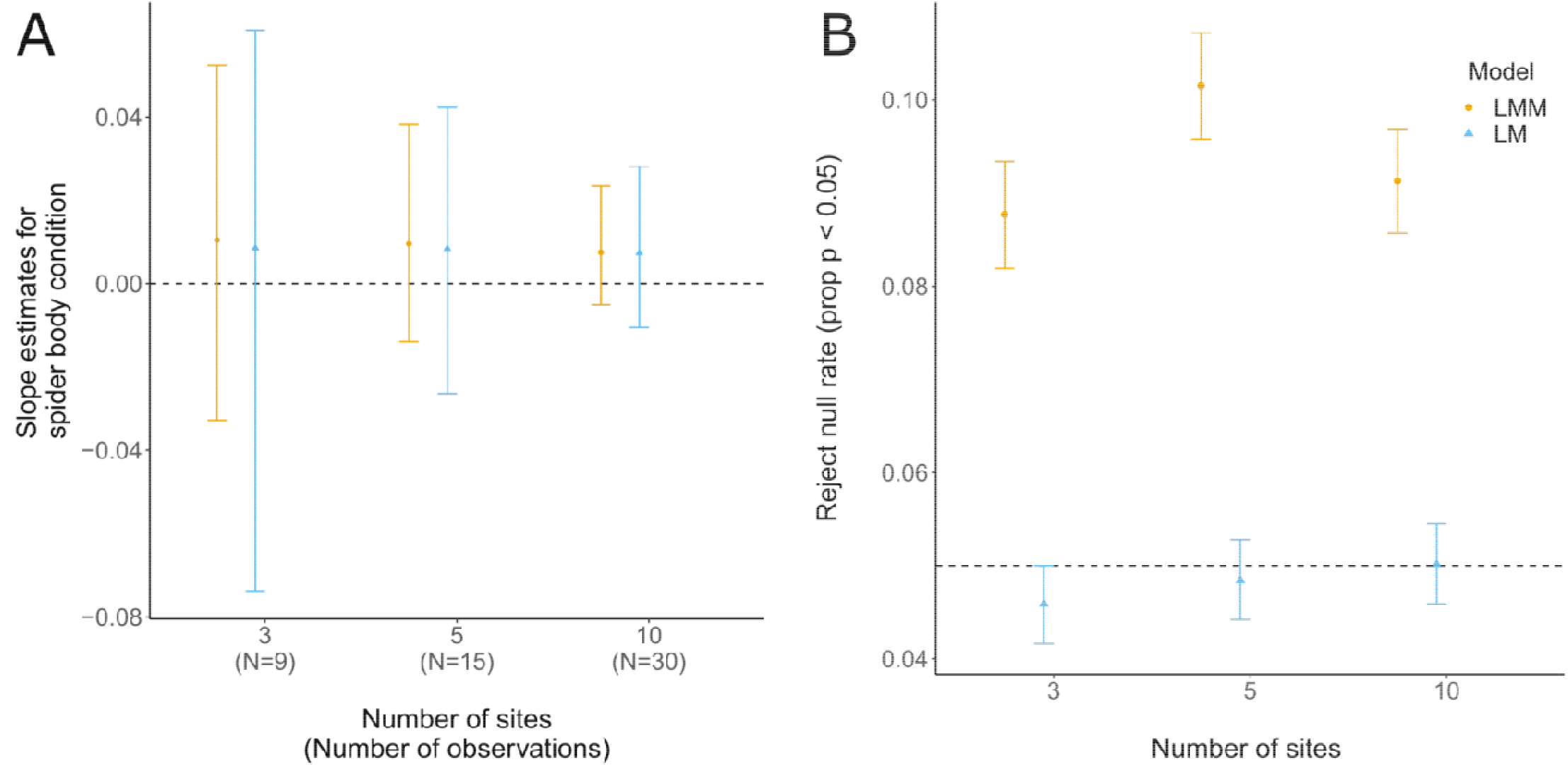
Analyses for real dataset (spiders in noise case study). (A): Fixed effects model estimates do not differ strongly across number of sites or model types, but estimates from linear models are slightly pulled toward zero. Each point in is the mean estimate for 10,000 models (and datasets), whereas error bars are 95% confidence intervals. N = the number of observations (i.e. number of rows) in each dataset. Dashed line at zero indicates the hypothesis that there is no effect of sound pressure level on spider body condition. As the sample size and the number of sites increases, our certainty around the parameter estimates increases. This is likely due to sample size rather than the number of sites (see Figure 3). Estimates from linear models (LM) are slightly more uncertain than estimates from linear mixed-effects models (LMM). (B): The probability for rejecting the null (*p* < 0.05) is higher for LMMs than LMs, suggesting that LMs are more conservative in this scenario. The probability of rejecting the null model for LMs overlaps alpha = 0.05, suggesting there is no effect of sound pressure level on spider body condition. Of course we do not know if rejecting the null model is “truth” here, since these are real biological data (see Figure 3 for simulated data). The probability of rejecting the null model was calculated as the proportion of models (n = 10,000) in which a ‘significant’ p value of ≤ 0.05 was obtained for the fixed effect parameter estimate; each point represents this proportion. To generate error bars as 95% confidence intervals, I used bootstrapping to replicate this process 1,000 times (see methods).

Output from 10,000 LMs suggests that the null hypothesis (no effect) would be rejected by a proportion of approximately 0.05 (Figure 4B), which is around the typical type-I error alpha level (i.e. suggesting that sound pressure level is not a significant predictor of spider body condition). LMMs, on the other hand, experienced null hypothesis rejection rates at nearly double that of their LM counterparts, which is consistent with the reduced uncertainty around the LMM estimates in Figure 4A. The rate of rejecting the null hypothesis, however, did not appear to be related to the number of sites (and thus total observations).

## Discussion

The work presented here demonstrates that i) fixed effects estimates are not more biased when the levels of an accompanying random effect have fewer than five (n < 5) levels, but population-level (random effects) variance estimates are and ii) type-I error rates are not increased by using linear models (LM) instead of linear mixed-effects models (LMM).

Fixed effects parameter estimation does not appear to be strongly influenced by, nor biased by, the number of levels of random effects terms. Instead, uncertainty in those estimates is much more strongly influenced by sample size. While this pattern may appear to contradict the decreased uncertainty (with more random effects levels) around beta estimates in Figure 2 of Harrison (2015) and in Figure 4 of the current work, this instead is due to differences in the way that sample size relates to the number of random effects levels. Harrison (2015) coded each random effect level to be associated with a fixed number of observations (N=20), such that each additional random effect level yielded an increased sample size – as is also the case in the spider case study presented here. However, in the simulations here (Figure 1), sample size (i.e. number of observations) has been separated from the number of random effects terms (e.g. sites or individuals).

Despite these differences in coding, the estimation of random effects terms (μ and σ) in the simulations here suggest consistent patterns with Harrison (2015) in that variance (σ) is more biased with fewer levels. This seems to support previous suggestions and simulations suggesting that few levels of random effects terms can make estimation of population-level variance difficult (Gelman & Hill, 2006; Harrison, 2015; Kéry & Royle, 2015; Harrison et al., 2018), but the cutoff at five random effects levels appears quite arbitrary. The combination of these results suggest that using fewer than five levels of random effects is acceptable when one is only interested in estimating fixed effects parameters (i.e. predictors, independent variables); in other words, when inference about the variance of random effects terms (e.g. sites, individuals, populations) is not of direct interest, but instead are used to group non-independent data. In these cases, however, caution should be taken in reporting the variance estimates for such population-level parameters – as this information can later be taken out of context of the question at hand and may result in the propagation of biased estimates.

Those following the “less than five levels” guideline typically drop random effects from analyses, turning a LMM into a LM. In both the simulations and the case study, LMMs and LMs did not appear to give drastically different parameter estimates for fixed effects. In the spider case study, LMMs gave more consistent results, leading to increased parameter certainty when compared to LMs, which was also reflected in a higher probability of ‘significant’ p values (around 10% of the models), when compared to LMs (around 5% of the models, which is to be expected to due chance). That is, results from LMs here suggest that there is no significant effect of the predictor (sound pressure level) on the response (body condition). While misspecified mixed-effects models can be overconfident in their estimates (Schielzeth & Forstmeier, 2009), in this case we do not know what ‘truth’ is. That is, we do not know if we *should* be rejecting the null hypothesis since the biological data here are real. However, this highlights that the p values might not always be very informative. For both model types in the case study (LMM vs LM), the magnitude of the parameter estimates (i.e. effect sizes) were consistently small. Interpreting the estimates directly will likely lead to a more consistent understanding of the results, rather than focusing on whether p values pass an arbitrary threshold.

The use, and abuse, of p values, in general, is highly debated and controversial (Yoccoz, 1991; Schervish, 1996; Wagenmakers, 2007; Murdoch, Tsai & Adcock, 2008; Gelman, 2013; Murtaugh, 2014; Leek & Peng, 2015; Greenland et al., 2016; Ho et al., 2019), but is further complicated by mixed-effects models (Luke, 2017). Douglas Bates, and other authors of the R package ‘lme4’ (Bates et al., 2007) which I use here, does not include a p value ‘baked in’ to the output for a reason (in short how one should calculate this is not straightforward or as obvious as it sounds). While my personal approach is generally to rely more on probabilistic (i.e. Bayesian) approaches or effect sizes and ignore p values wherever possible, I use p values here because they are still widely used across ecology and evolutionary studies and by fish and wildlife managers. The R package ‘lmerTest’ (Kuznetsova, Brockhoff & Christensen, 2017) allows many users to get around this *apparent* limitation of ‘lme4’ by calculating p values as if they were a part of the original package. Despite my recommendation to focus more on effect size and other model outputs in analyses of real data, understanding the consequences of type-1 errors in terms of p values is relevant so long as ecologists continue to use them.

Interestingly, type-I errors were not more likely in any of the LMs of simulated data. This possibly suggests that misspecified linear models (theoretically missing a random effect) are relatively robust to this omission – at least in some simple cases such as the scenarios presented here. While this perhaps alleviates some concern over inflated type-I errors due to pseudoreplication while ignoring the grouped nature of repeat-measures studies and non-independent data (Arnqvist, 2020), this should not be taken as evidence to purposefully omit random effects when such a structure is appropriate. Instead, it warrants future investigation and further simulation studies with more thorough scenarios (especially with varying degrees of random effect variance) and more complex data structures (e.g. including correlations and link functions).

Often researchers (sometimes nudged by peer-reviewers) cite this guideline of needing 5 levels before random effects inclusion as a reason why they were unable to use a mixed-effects model (Bain, Johnson & Jones, 2019; Bussmann & Burkhardt-Holm, 2020; Evans & Gawlik, 2020; Gomes & Goerlitz, 2020; Zhao, Johnson-Bice & Roth, 2021). Although there is confusion over this recommendation, as some opt to use mixed-effects models despite this suggestion (Latta et al., 2018; Fugère, Lostchuck & Chapman, 2020; Gomes, Appel & Barber, 2020; Allen et al., 2021), likely because of the numerous advantages that mixed-effects models offer (Bolker, 2008; Kéry & Royle, 2015; Harrison et al., 2018), or fear of the consequences of pseudoreplication (although this can easily occur in mixed-effects models as well: Schielzeth & Forstmeier, 2009; Arnqvist, 2020). The trend to automatically follow this rule is likely exacerbated by the fact that authors or peer-reviewers can easily point out that this rule exists (Gelman & Hill, 2006; Harrison, 2015; Kéry & Royle, 2015; Harrison et al., 2018; Arnqvist, 2020), but may find it more difficult or time-consuming to make a nuanced argument against following such a rapidly growing rule. Hopefully the results presented here will challenge that view, and allow the fitting of random effects when inference is not being made for the random effects. More importantly, I hope it sparks further conversation and debate over this issue. Given the widespread accessibility of GLMMs, future simulation studies and further assessments of these statistical methods are necessary to understand the consequences of both violating and blindly following methodological rules.

## Acknowledgements

I would like to thank Xavier Harrison for inspiration, Trevor Caughlin for aiding in my understanding of simulations, the National Science Foundation (NSF) for funding (GRFP 2018268606), and Boise State University and the Hatfield Marine Science Center at Oregon State University for general and logistical support.

---

**Figure S1:**
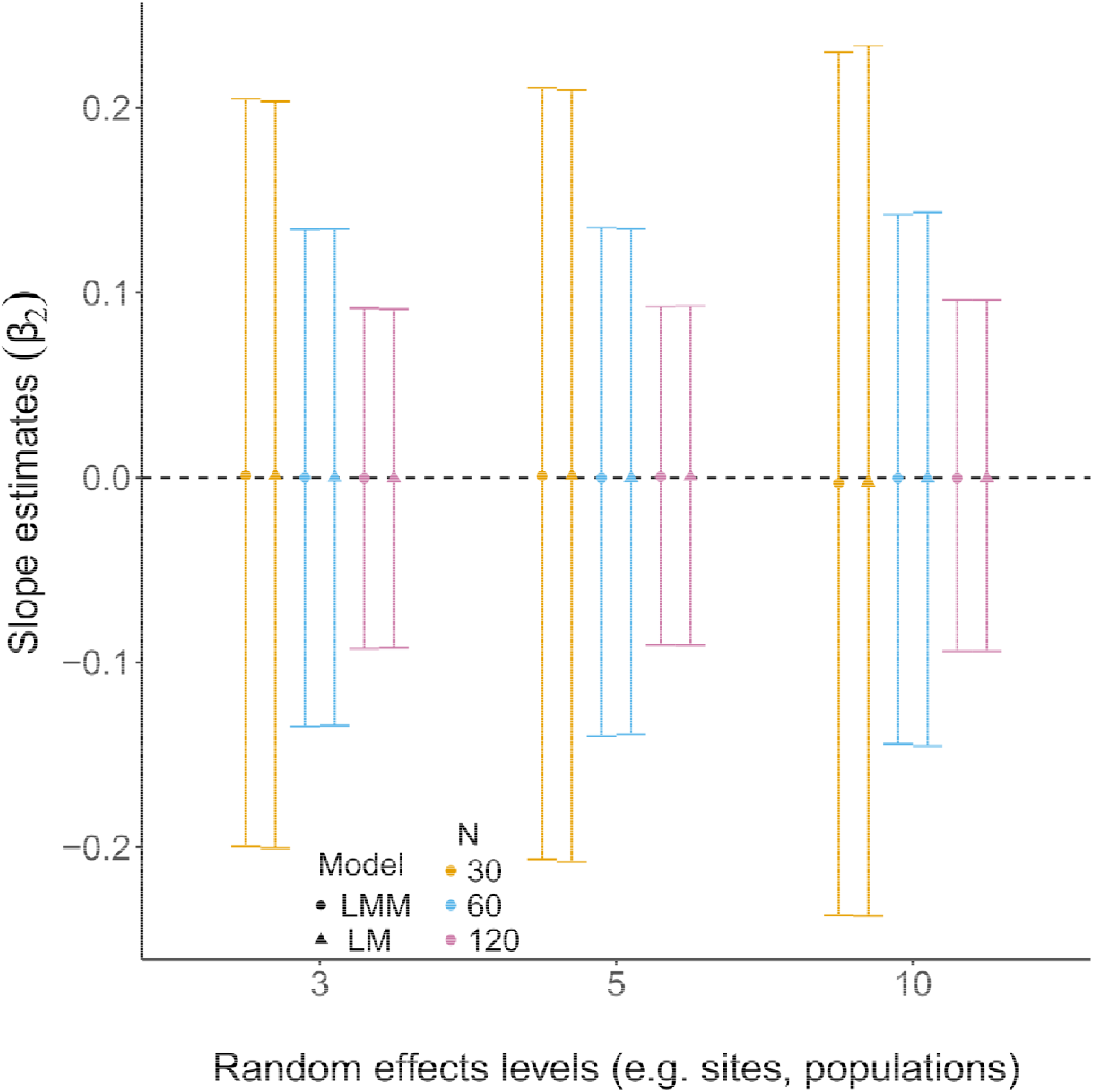
A comparison of LMM and LM parameter estimates. Each point is the mean estimate for 10,000 models (and datasets), whereas error bars are 95% confidence intervals. N = the number of observations (i.e. number of rows) in each data set. Dashed lines indicate the true value. In all scenarios the bias in parameter estimates are negligible. As the sample size increases, our certainty around the parameter estimates (β) increases, but the number of random effects in the data set has a relatively minor effect on estimating β. When sample sizes (N) are low, parameter uncertainty increases with increasing levels of random effects (assuming a consistent N). With linear models (triangles), random effects are not fitted in the model, but they were still present during the data-generating process. Here, linear models do just as well as linear mixed-effects models at resurrecting true parameter values.

## References

Allen LC, Hristov NI, Rubin JJ, Lightsey JT, Barber JR. 2021. Noise distracts foraging bats. Proceedings of the Royal Society B 288:20202689.

Arnqvist G. 2020. Mixed models offer no freedom from degrees of freedom. Trends in ecology & evolution 35:329–335.

Bain GC, Johnson CN, Jones ME. 2019. Chronic stress in superb fairy-wrens occupying remnant woodlands: Are noisy miners to blame? Austral Ecology 44:1139–1149.

Barr DJ, Levy R, Scheepers C, Tily HJ. 2013. Random effects structure for confirmatory hypothesis testing: Keep it maximal. Journal of memory and language 68:255–278.

Bates D, Sarkar D, Bates MD, Matrix L. 2007. The lme4 package. R package version 2.

Bolker BM. 2008. Ecological models and data in R. Princeton University Press.

Bolker BM, Brooks ME, Clark CJ, Geange SW, Poulsen JR, Stevens MHH, White J-SS. 2009. Generalized linear mixed models: a practical guide for ecology and evolution. Trends in ecology & evolution 24:127–135.

Bürkner P-C. 2017. brms: An R package for Bayesian multilevel models using Stan. Journal of statistical software 80:1–28.

Bussmann K, Burkhardt-Holm P. 2020. Round gobies in the third dimension-use of vertical walls as habitat enables vector contact in a bottom-dwelling invasive fish. Aquatic Invasions 15:683–699.

Carpenter B, Gelman A, Hoffman MD, Lee D, Goodrich B, Betancourt M, Brubaker MA, Guo J, Li P, Riddell A. 2017. Stan: a probabilistic programming language. Grantee Submission 76:1–32.

Evans BA, Gawlik DE. 2020. Urban food subsidies reduce natural food limitations and reproductive costs for a wetland bird. Scientific reports 10:1–12.

Fugère V, Lostchuck E, Chapman LJ. 2020. Litter decomposition in Afrotropical streams: Effects of land use, home-field advantage, and terrestrial herbivory. Freshwater Science 39:497–507.

Gabry J, Goodrich B. 2016. rstanarm: Bayesian applied regression modeling via Stan. R package version 2.10. 0.

Gelman A. 2008. Scaling regression inputs by dividing by two standard deviations. Statistics in medicine 27:2865–2873.

Gelman A. 2013. Commentary: P values and statistical practice. Epidemiology 24:69–72.

Gelman A, Hill J. 2006. Data analysis using regression and multilevel/hierarchical models. Cambridge university press.

Gomes DGE. 2020. Orb-weaving spiders are fewer but larger and catch more prey in lit bridge panels from a natural artificial light experiment. PeerJ 8:e8808.

Gomes DG, Appel G, Barber JR. 2020. Time of night and moonlight structure vertical space use by insectivorous bats in a Neotropical rainforest: an acoustic monitoring study. PeerJ 8:e10591.

Gomes DGE, Goerlitz HR. 2020. Individual differences show that only some bats can cope with noise-induced masking and distraction. PeerJ 8:e10551.

Gomes DGE, Hesselberg T, Barber JR. 2021. Phantom river noise alters orb-weaving spider abundance, web size, and prey capture. Functional Ecology 35:717–726.

Greenland S, Senn SJ, Rothman KJ, Carlin JB, Poole C, Goodman SN, Altman DG. 2016. Statistical tests, P values, confidence intervals, and power: a guide to misinterpretations. European journal of epidemiology 31:337–350.

Harrison XA. 2015. A comparison of observation-level random effect and Beta-Binomial models for modelling overdispersion in Binomial data in ecology & evolution. PeerJ 3:e1114.

Harrison XA, Donaldson L, Correa-Cano ME, Evans J, Fisher DN, Goodwin CE, Robinson BS, Hodgson DJ, Inger R. 2018. A brief introduction to mixed effects modelling and multi-model inference in ecology. PeerJ 6:e4794.

Ho J, Tumkaya T, Aryal S, Choi H, Claridge-Chang A. 2019. Moving beyond P values: data analysis with estimation graphics. Nature methods 16:565–566.

Hurlbert SH. 1984. Pseudoreplication and the design of ecological field experiments. Ecological monographs 54:187–211.

Jakob EM, Marshall SD, Uetz GW. 1996. Estimating fitness: a comparison of body condition indices. Oikos:61–67.

Kass RE, Caffo BS, Davidian M, Meng X-L, Yu B, Reid N. 2016. Ten simple rules for effective statistical practice. Public Library of Science.

Kéry M, Royle JA. 2015. Applied Hierarchical Modeling in Ecology: Analysis of distribution, abundance and species richness in R and BUGS: Volume 1: Prelude and Static Models. Academic Press.

Kuznetsova A, Brockhoff PB, Christensen RH. 2017. lmerTest package: tests in linear mixed effects models. Journal of statistical software 82:1–26.

Latta SC, Brouwer NL, Mejía DA, Paulino MM. 2018. Avian community characteristics and demographics reveal how conservation value of regenerating tropical dry forest changes with forest age. PeerJ 6:e5217.

Leek JT, Peng RD. 2015. Statistics: P values are just the tip of the iceberg. Nature News 520:612.

Low-Décarie E, Chivers C, Granados M. 2014. Rising complexity and falling explanatory power in ecology. Frontiers in Ecology and the Environment 12:412–418.

Luke SG. 2017. Evaluating significance in linear mixed-effects models in R. Behavior research methods 49:1494–1502.

Magnusson A, Skaug H, Nielsen A, Berg C, Kristensen K, Maechler M, van Bentham K, Bolker B, Brooks M, Brooks MM. 2017. Package ‘glmmTMB.’ R Package Version 0.2. 0.

McElreath R. 2020. Statistical rethinking: A Bayesian course with examples in R and Stan. CRC press.

Murdoch DJ, Tsai Y-L, Adcock J. 2008. P-values are random variables. The American Statistician 62:242–245.

Murtaugh PA. 2014. In defense of P values. Ecology 95:611–617.

Patil A, Huard D, Fonnesbeck CJ. 2010. PyMC: Bayesian stochastic modelling in Python. Journal of statistical software 35:1.

Powell LA, Gale GA. 2015. Estimation of Parameters for Animal Populations. Caught Napping Publications, Lincoln, NE.

R Core Team. 2017. R: A language and environment for statistical computing. Vienna, Austria: R Foundation for Statistical Computing.

Rue H, Riebler A, Sørbye SH, Illian JB, Simpson DP, Lindgren FK. 2017. Bayesian computing with INLA: a review. Annual Review of Statistics and Its Application 4:395–421.

Salvatier J, Wiecki TV, Fonnesbeck C. 2016. Probabilistic programming in Python using PyMC3. PeerJ Computer Science 2:e55.

Schervish MJ. 1996. P values: what they are and what they are not. The American Statistician 50:203–206.

Schielzeth H, Forstmeier W. 2009. Conclusions beyond support: overconfident estimates in mixed models. Behavioral ecology 20:416–420.

Silk MJ, Harrison XA, Hodgson DJ. 2020. Perils and pitfalls of mixed-effects regression models in biology. PeerJ 8:e9522.

Wagenmakers E-J. 2007. A practical solution to the pervasive problems of p values. Psychonomic bulletin & review 14:779–804.

Wickham H. 2011. ggplot2. Wiley Interdisciplinary Reviews: Computational Statistics 3:180–185.

Yoccoz NG. 1991. Use, overuse, and misuse of significance tests in evolutionary biology and ecology. Bulletin of the Ecological Society of America 72:106–111.

Zhao S-T, Johnson-Bice SM, Roth JD. 2021. Foxes facilitate other wildlife through ecosystem engineering activities on the Arctic tundra. bioRxiv.

Zuur AF, Ieno EN. 2016. A protocol for conducting and presenting results of regression-type analyses. Methods in Ecology and Evolution 7:636–645.

Zuur AF, Ieno EN, Elphick CS. 2010. A protocol for data exploration to avoid common statistical problems. Methods in ecology and evolution 1:3–14.

